# IgA plasma cells co-secrete monomeric and dimeric IgA

**DOI:** 10.64898/2026.07.03.736325

**Authors:** Jana Thomas, Klaus Eyer, Jens Wittner, Tim Rollenske, Edith Roth, Wei Xiang, Wolfgang Schuh, Hans-Martin Jäck, Dirk Mielenz, Sebastian R. Schulz

**Affiliations:** Department of Translational Immunology, Nikolaus-Fiebiger Center of Molecular Medicine, Universitätsklinikum Erlangen, Friedrich-Alexander-Universität Erlangen-Nürnberg, Glückstr. 6, 91054 Erlangen, Germany; Department of Biomedicine, Aarhus University, Wilhelm Meyers Allé 4, 8000 Aarhus, Denmark; Division of Molecular Immunology, Department of Internal Medicine 3, Nikolaus-Fiebiger Center of Molecular Medicine, Universitätsklinikum Erlangen, Friedrich-Alexander-Universität Erlangen-Nürnberg, Glückstr. 6, 91054 Erlangen, Germany; University of Bonn, University Hospital Bonn, Institute of Molecular Medicine and Experimental Immunology, Bonn, Germany; Translational Radiobiology, Department of Radiation Oncology, Universitätsklinikum Erlangen, Friedrich Alexander Universität Erlangen-Nürnberg, Erlangen, Germany; Department of Molecular Neurology, Department Kopfklinik, Universitätsklinikum Erlangen, Friedrich-Alexander-Universität Erlangen-Nürnberg, Schwabachanlage 6, 91054 Erlangen, Germany

**Keywords:** Plasma cell, IgA, dimeric IgA, poly Ig receptor

## Abstract

Dimeric immunoglobulin A (dIgA) is generated from IgA monomers (mIgA) via JCHAIN-dependent polymerization. DIgA is transported across epithelial barriers by the poly Ig receptor (PIGR) and confers mucosal protection, while serum contains substantial amounts of IgA monomers. Distinct plasma cell subsets have been proposed to produce either monomeric or dimeric IgA, with bone marrow plasma cells as a primary source of mIgA. Here, we addressed whether IgA plasma cell populations segregate based on mIgA or dIgA production. Flow cytometric analysis of antibody-secreting cells from bone marrow, lymphoid and mucosal tissues revealed universal intracellular JCHAIN expression across isotypes and failed to identify a discrete JCHAIN-negative IgA plasma cell population. To detect polymeric IgA, we generated a recombinant soluble PIGR that selectively bound JCHAIN-containing dIgA in Western blot, ELISA, and flow cytometry. Soluble PIGR binding was detected in all IgA plasma cells irrespective of tissue origin, arguing against a dedicated mIgA-producing plasma cell subset incapable of dIgA formation. Ex vivo cultures and single-cell DropMap secretion assays demonstrated that bone marrow and lamina propria IgA antibody-secreting cells co-secrete mIgA and dIgA. These findings suggest that dIgA assembly and secretion are general properties of IgA plasma cells and disfavor a dedicated mIgA-producing population.

**Highlights:**

- All plasma cells express JCHAIN protein
- Recombinant poly Ig receptor detects dimeric IgA in all IgA plasma cells
- Bone marrow plasma cells secrete dimeric IgA
- IgA plasma cells co-secrete mono- and dimeric IgA

## Introduction

Immunoglobulin A (IgA) is the most abundant Ig isotype in humans and the second most abundant isotype in serum. It is the dominant Ig isotype in mucosal cavities and breast milk, safeguarding mucosal surfaces against microbial invasion and maintaining immune homeostasis at the mucosal barrier (1,2). The formation of IgA involves the covalent assembly of two IgA heavy chains and two light chains into monomeric IgA (mIgA). However, plasma cells also produce the joining chain (JCHAIN), which can form disulfide bridges with IgA and IgM monomers, creating IgA dimers or IgM pentamers (3–5). Yet, ∼ 80% of human serum IgA is monomeric, while mucosal IgA is dimeric. Notably, dimerization of IgA is a prerequisite for mucosal transport across the lining epithelium, because only JCHAIN containing IgA can bind to the polymeric Ig receptor (PIGR) expressed on the basolateral surface of epithelial cells (6). The PIGR mediates transcytosis of dimeric IgA (dIgA) across the epithelium. During this process, the PIGR undergoes proteolytic cleavage, releasing the secretory component (SC) bound to dIgA, forming secretory IgA (sIgA) (6). sIgA dominates the mucosal surfaces while blood contains only residual amounts of dIgA, as it is rapidly absorbed into the mucosa (7). Thus, monomeric IgA is retained in the blood, where, in humans, it can engage with the Fcα receptor I expressed on immune cells, such as neutrophils, monocytes, and certain dendritic cell subsets (8,9). Therefore, IgA dimerization is key to its spatial function: sIgA, due to its increased avidity, provides strong protection against pathogens, often leading to immunologically silent removal before they can cross the mucosal barrier (10,11). In the case of pathogen breakthrough, however, systemic IgA can then potently activate immune cells and elicit strong inflammatory reactions (12). The complex functions of IgA are reflected by the fact that selective IgA deficiency, albeit often asymptomatic, can lead to a variety of clinical symptoms, ranging from sinopulmonary infections and allergy to autoimmunity (13). While dIgA exhibits superior virus-neutralizing activity compared to mIgA (14–16), it is also a key feature of autoreactive antibody responses (17). To understand and potentially regulate immune responses, it is key to identify the mechanisms of IgA dimerization. Conceptually, the regulation of IgA dimerization may occur in a cell-intrinsic manner via regulation of JCHAIN expression or chaperones, such as marginal zone B and B 1 cell specific protein 1 (MZB1) (5,18). Alternatively, dedicated mIgA- or dIgA-producing lineages might exist, with dIgA-producing plasma cells thought to be aligned along mucosal membranes (10,19). In our study, we evaluate both models in mice. To this end, we established a soluble PIGR (sPIGR) reagent to detect fully assembled dIgA by ELISA, Western blot and flow cytometry. We compared IgA and JCHAIN protein abundance with PIGR-binding. Thereby, we found that all IgA plasma cells, regardless of their anatomical origin, produce JCHAIN and dimeric IgA, which disfavors the lineage model of a distinct plasma cell population responsible for mIgA. We propose a model in which the regulation of mIgA and dIgA production is controlled cell-intrinsically.

## Methods

### Mice and sample collection

C57BL/6N mice were purchased from Janvier (Le Genest Saint Isle, France, stock no. C57BL/6NRj) and maintained under specific pathogen-free conditions at the Nikolaus-Fiebiger Center animal facility, Friedrich-Alexander University Erlangen–Nürnberg, Germany. Mice were used for experiments at 15-25 weeks of age. Animals were euthanized by carbon dioxide inhalation prior to tissue collection. All animal procedures were performed in accordance with institutional and national guidelines and approved by the relevant local authority (TS 05/07, Amt für Veterinärwesen und gesundheitlichen Verbraucherschutz der Stadt Erlangen, Erlangen, Germany; Regierung von Unterfranken, Würzburg, Germany). For DropMap experiments, female C57BL/6N mice aged 7-9 weeks were used. These animal experiments were approved by the Danish Animal Experiments Inspectorate (Dyreforsøgstilsynet; approval no. 2025-20054), granted to Klaus Eyer.

Fecal samples were homogenized in PBS (Gibco Life Technologies, Cat.:14190-094) at a ratio of 100µL per mg material. Samples were then centrifuged (13,000 rpm, 10min, 4°C), supernatants were collected and stored at −20°C until analysis. To isolate murine saliva, the mice were allowed to chew on the oral swab. The saliva was then recovered from the swab by centrifugation and stored at −20 °C. Blood was obtained by cardiac puncture and collected in Microtainer blood collection tubes (BD, 365968). Serum was prepared according to the manufacturer’s instructions.

### Isolation of colon lamina propria cells

Colon was isolated, residual fat tissue and Peyer’s patches removed. The tissue was longitudinally opened, and thoroughly washed with PBS (Gibco Life Technologies, Cat.:14190-094) supplemented with 2% heat-inactivated FCS (Gibco Life Technologies, Cat.:10270-106). Tissue was cut into approximately 0.5 cm pieces and incubated twice for 15 min in R10 medium consisting of RPMI 1640 (Gibco Life Technologies, Cat.:31870-025), supplemented with 10% FCS, 2 mM L-glutamine (Gibco Life Technologies, Cat.:25030-024), 1 mM sodium pyruvate (Gibco Life Technologies, Cat.:11360-039), 50 U/ml penicillin, 50 µg/ml streptomycin (Gibco Life Technologies, Cat.:15140-122), 50 µM β-mercaptoethanol (Gibco Life Technologies, Cat.:31350-010), with additional 25 mM EDTA (Invitrogen, 15575-038), at 37°C under continuous rotation at 230rpm. Following washing with RPMI 1640, the tissue was minced with scissors and transferred to gentleMACS C Tubes (Miltenyi Biotec, 130-093-237) containing 5 ml pre-warmed digestion solution prepared from the Lamina Propria Dissociation Kit (Miltenyi Biotec, 130-097-410; 100 µl Enzyme D, 50 µl Enzyme R, and 12.5 µl Enzyme A). The sample was then processed using a gentleMACS Octo Dissociator (Miltenyi Biotec, 130-134-029; program 37C_m_LPDK_1). The digested suspension was filtered through 70µm cell strainers (Corning, 431751), washed with RPMI 1640, centrifuged (350xg, 4°C) for 7 min, and resuspended in R10 medium for cell counting.

### Flow cytometry

Single-cell suspensions of cLP were isolated as described above and additional tissues were processed as previously described (25). Samples were washed (1400rpm, 7min, 4°C) in PBS (Gibco Life Technologies, Cat.:14190-094) containing 2% fetal FCS (Gibco Life Technologies, Cat.:10270-106) and stained with fluorochrome-conjugated antibodies for 20 min on ice in the dark. Following washing, cells were fixed with Reagent A for 15 minutes at RT in the dark and afterwards permeabilized using Reagent B (Nordic MuBio Kit, GAS-002-1-CE/IVD) for 20 minutes at RT in the dark. Intracellular staining was performed in permeabilization buffer. Samples were acquired on a Gallios flow cytometer (Beckman Coulter) and analysed using Kaluza software (Beckman Coulter). Compensation matrices were generated using single-stained controls. Fluorescence-minus-one (PBS + anti-rabbit IgG antibodies) and FLAG-tag controls (RBD + anti-FLAG antibodies) were used to define JCHAIN and sPIGR gates. The following antibodies were used for flow cytometry stainings: anti-CD138-PE.Cy7 (BioLegend, Clone 281-2, dilution 1:1500), anti-TACI-PE (eBioscience, Clone eBio8F10-3, dilution 1:400), anti-ENPP1-BV421 (BioLegend, Clone YE1/19.1, dilution 1:800), anti-CD98-PE (BioLegend, Clone RL388, dilution 1:2000), anti-JCHAIN (Invitrogen, Clone SP105, dilution 1:100), recombinant sPIGR (5 µg/ml), anti-IgM-APC.Cy7 (BioLegend, Clone RMM-1, dilution 1:100), anti-IgA-FITC (Southern Biotech, Cat.: 1040-02, dilution 1:1000) and for secondary detection anti-rabbit IgG (H+L)- AF647 (Jackson, Cat.: 111-605-144, dilution 1:500), anti-rabbit IgG-PE (Biolegend, Cat.: Poly4064, dilution 1:500), anti-FLAG (DYK)-APC (BioLegend, Clone L5, dilution 1:200), Strep-Tactin XT-Dy649 (IBA, 2-1568-050, dilution 1:400)

### Generation of J558 Jchain-deficient cell lines

Two sgRNAs targeting exon 2 of the mouse Jchain locus (5′-UGUGUACCCGAGUUACCUCU-3′ and 5′-GCAUUUGUUGUCAGCAAGAA-3′) were designed using the Synthego sgRNA design tool and purchased from Synthego. The two sgRNAs were pooled and assembled with recombinant Cas9 protein (Synthego) at equimolar ratios to generate ribonucleoprotein complexes. J558 cells were transfected using the Neon electroporation system (Thermo Fisher Scientific) with 10 µl Neon tips and the following parameters: 1300 V, two pulses, and 20 ms pulse width. After recovery, single cells were FACS-sorted into 96-well plates and expanded for 12 days. Clones were screened by intracellular flow cytometry using an anti-JCHAIN antibody (Invitrogen, clone SP105). Selected JCHAIN-deficient clones and JCHAIN-positive control clones were further expanded in R10 medium.

### Plasmid construction and HEK293T transfection

A codon-optimized expression construct encoding mouse JCHAIN (Reference Sequence: NP_690052.2; amino acids 1-159) with a C-terminal FLAG tag was synthesized and cloned into a pcDNA3.1 backbone by GenScript. The complete J558 mouse IgA heavy-chain coding sequence was amplified from J558 cDNA and assembled into a pcDNA3.1 backbone using NEBuilder HiFi DNA Assembly Master Mix (NEB, Cat.: E2621S). HEK293T cells were transiently transfected using polyethyleneimine, as described previously (45) and analyzed by intracellular flow cytometry 24 h after transfection

### Expression and purification of soluble PIGR

The coding sequence corresponding to the extracellular domain of mouse PIGR (Reference Sequence NP_035212.2, amino acids 21-611) was cloned into the pcDNA3 expression vector under the control of the CMV promoter, with an N-terminal mouse Igkv3-10 signal peptide (METDTLLLWVLLLWVPGSTG). A Twin-Strep-tag followed by a 3×FLAG tag was added at the C-terminus. 293F cells (Gibco, Cat.: R79007) were transfected with the sPIGR expression plasmid using polyethylenimine (Polysciences, Cat.: 24765) and maintained in serum-free FreeStyle 293 Expression Medium (Gibco, Cat.: 12338026) at 37°C/8% CO_2_ at 125rpm in an orbital shaker incubator. After 72h, culture supernatants were harvested and purified using an ÄKTA Pure 25 L system (GE Healthcare) and a StrepTrap XT 1ml column (GE Healthcare, Cat.: GE29401317). The column was equilibrated with buffer W (iba, Cat.: 2-1003-100) and protein eluted with buffer BXT (iba, Cat.: 2-1042-025). The eluted protein was dialyzed against PBS (Gibco Life Technologies, Cat.:14190-094). Purity and integrity were verified by Coomassie staining of 10% SDS-PAGE. Gel electrophoresis was performed as described for Western blotting, under reducing and non-reducing conditions. Gels were stained for 10 min with Coomassie staining solution (0.1% Coomassie Brilliant Blue R-250 (Sigma-Aldrich, Cat.:27816), 50% methanol (Roth, Cat.: T909.1), 10% acetic acid (Roth, Cat.: 6755.1) in dH₂O), followed by destaining in 40% methanol and 10% acetic acid in dH₂O.

### Western Blot

Murine saliva, feces, serum and cell culture supernatants were analyzed under non-reducing Western blot conditions for monomeric and dimeric IgA. Samples were mixed with 5x non-reducing loading dye in a ratio 1:4 (400 mM Tris-HCl (Roth, Cat.: 4855.3) pH 6.8, 20% glycerol (Roth, Cat.: 3783.1), 10% SDS (Roth, Cat.: 2326.2), 0.01% Bromophenol blue (Merck, Cat.: 108122)) and heated to 60°C for 5 minutes. For reducing conditions, 100 mM Di-thio-threitol (Gibco, 31350-010) were added, followed by incubation at 95°C for 5 minutes. Samples were analyzed at the following final dilutions: supernatant 1:1.25, feces 1:25, saliva 1:25 and sera 1:1000. Cell culture supernatants were generated by incubating 2 x 10⁵ J558 cells in 1 mL R10 medium for 3 days. All samples were loaded onto gels with 6 % acrylamide (non-reducing) or 12.5 % acrylamide (reducing), together with PageRuler Prestained Protein Ladder (Thermofisher, Cat.: 26617). Electrophoresis was performed at 60-70mA (max. 400V, 60W) per gel for 3-4h. Separated proteins were transferred onto nitrocellulose membranes (Amersham, Cat.: GE10600001) by semi-dry blotting at 400mA (max. 50V, 10W) for 45min. The membrane was blocked in TBS-T buffer (150 mM NaCl (Roth, Cat.: 0962.2)), 20 mM Tris/HCl (Roth, Cat.: 2449.3), pH 7.4 containing 0.1% Tween-20 (Sigma-Aldrich, Cat.: P1379)) with 5 % milk (Roth, Cat.: T145.3) for at least 1h and washed four times with TBS-T (5-10min each time). Proteins of interest were detected by incubating the membrane with the respective antibody for 1h at RT or overnight at 4°C. For IgA detection, a 1:10,000 dilution of HRP-conjugated anti-mouse IgA antibody (Southern Biotech, Cat.: 1040-05) in TBS-T with 5% milk was used. JCHAIN was detected using the primary anti-human/mouse JCHAIN antibody (Invitrogen, Cat.: MA5-16419) at 30 ng/ml in antibody buffer (3% BSA (Roth, 8076.4), 0.1% Tween 20, 0.1% Sodium Azide (Roth, Cat.: 4221.1) in PBS), followed by a 1:10,000 dilution of HRP-conjugated anti-rabbit IgG antibody (Southern Biotech, Cat.: 1030-05) in TBS-T with 5% milk. For detection of sPIGR-binding proteins, membranes were incubated with 1.6 µg/ml sPIGR in antibody buffer, followed by a 1:2,000 dilution of HRP-conjugated anti-FLAG antibody (Biolegend, Cat.: 637311) in TBS-T containing 5% milk. After washing, the membranes were incubated for 30 s with ECL solution (GeneTex, Cat.: GTX14698) and then exposed to an CL-Xposure films (Thermo Scientific, Cat.: 34089) for 10 s to 20 min.

### Culture of bulk-sorted IgA antibody-secreting cells

For cLP and bone marrow cultures, cell suspensions were prepared from C57BL/6 mice as described and pooled from 4 animals per sample. IgA ASCs (CD138+ TACI+ IgA+) were then isolated using a MoFlo Astrios cell sorter (Beckman Coulter) and sorted ASCs were cultured at a density of 250,000 cells per ml in R10 medium supplemented with 50ng/ml recombinant APRIL (AdipoGen Life Sciences, Cat.: AG-40B-0089-3010) and 25mM HEPES (Gibco Life Technologies, Cat.: 15630080). Culture supernatants were collected after incubation for 24h at 37°C and 5% CO₂ and used for subsequent ELISA analysis.

### ELISA

Total IgA and dimeric IgA concentrations were quantified by ELISA. For calibration of the sPIGR-based assay, defined mixtures of recombinant monomeric and dimeric IgA were prepared (31), maintaining a constant total IgA concentration of 200 ng/ml. Pure recombinant dimeric IgA and an unconjugated IgA antibody (Sigma-Aldrich, clone TEPC15) were used as standards for dIgA and total IgA quantifications, respectively. Flat-bottom 96-well plates (Greiner Bio-One, Cat.: 655001) were coated overnight at 4 °C with 50 µl goat-anti-mouse IgA antibody (SouthernBiotech, Cat.: 1040-01, dilution 1:1000) in coating buffer (15 mM Na₂CO₃ (Supelco, Cat.: 1063921000), 35 mM NaHCO₃ (Supelco, Cat.: 1063291000) in dH2O). Plates were washed three times with washing buffer (PBS, 0.05% Tween-20 (Sigma-Aldrich, Cat.: P1379)) and blocked with 200 µl blocking buffer (PBS, 2% BSA (Roth, Cat.: 8076.4) overnight at 4°C. Sample and standard were loaded in duplicates onto the plates and were stepwise diluted 1:2 with blocking buffer. Following incubation for 1 h at RT, plates were washed three times with washing buffer. For total IgA detection, wells were incubated for 1 h with 50 µl HRP-conjugated goat-anti-mouse IgA antibody (SouthernBiotech, Cat.: 1040-05, 1:1000). For dIgA detection, plates were incubated with 50 µl recombinant sPIGR (5 µg/ml) for 1 h, washed, and subsequently incubated with 50 µl HRP-conjugated rat-anti-FLAG antibody (BioLegend, clone L5, dilution 1:2000) for 1h. After washing, colour reaction was induced by 50 µl TMB substrate (BD OptEIA Kit, Cat.: 555214). The reaction was stopped with additional 50 µl 0.5 M H₂SO₄ (Roth, Cat.: K027.2), and absorbance was measured at 450 nm using a Mini ELISA Plate Reader (BioLegend).

### DropMap analysis

For fabrication and set-up of microfluidic preparations – chips, chambers and solutions – we refer to (46). For the droplet-based measurement, the cells were sorted into ice-cold assay buffer consisting of RPMI-1640 without phenol red (Gibco Life Technologies, Cat.: 11835030), supplemented with 10% (v/v) inactivated fetal calf serum (Gibco Life Technologies, Cat.:10270-106), 1X penicillin–streptomycin (Gibco Life Technologies, Cat.:15140122), 10mM HEPES (Gibco Life Technologies, Cat.:15630056), 0.1% (v/v) Pluronic F-127 (Invitrogen, P6867), pH 7.2-4 with additional 0.5% (w/v) recombinant human serum albumin (Sigma-Aldrich, Cat.:A1653). Sorted cells were directly encapsulated post-sort using standard droplet microfluidics as described elsewhere (46). J558 wildtype and JCHAIN knock-out cells were collected by centrifugation, counted, and resuspended in a respective volume of assay buffer to achieve an encapsulation rate of 0.2-0.3 cells/droplet. For calibration and assay setup, we used recombinant murine IgA (dimer and monomer) (47). Paramagnetic nanoparticles (Streptavidin Plus, 300 nm, Ademtech) were used to assess the secretion of total and dIgA, and they were prepared as described elsewhere (46). For measuring primary cells from mice and calibration, we used CaptureSelect™ Biotin Anti-LC-kappa Mouse conjugate, whereas CaptureSelect™ Biotin Anti-LC-lambda Mouse conjugate was used to assess IgA secretion from J558 wildtype (clone E11) and JCHAIN knock-out cells (clone E1) . The IgA assay contained anti-IgA-FITC (Southern Biotech, Cat.: 1040-02) and anti-FLAG (DYKDDDDK)-AF647 (BioLegend, clone L5). Data was recorded and analyzed as previously described (46). For the different samples, an array of 10×10 images (cell lines) or 15x15 images (primary cells), corresponding to around 50,000-112,000 droplets, were acquired for 2 hours (cell lines) or overnight (primary cells). The images were analyzed using an updated custom Matlab script (Mathworks) (32). Values were exported to Excel (Microsoft), and the data further processed. The ratio between the mean fluorescence signal on the nanoparticles and the mean fluorescence signal of the whole droplet was determined, and only droplets containing signals corresponding to >10 nM total IgA (corresponding to a relocation signal of 1.4 in the FITC signal) were further selected. Their increase in ratios anti-IgA-FITC and anti-FLAG (DYKDDDDK)-AF647 was determined, and the final selected droplets were visually controlled and sorted for secretion or signals on cells.

## Statistical analysis

Quantification and statistical analyses were performed using GraphPad Prism version 10.6.1. Data distribution was assessed using the Shapiro–Wilk normality test. Comparisons between two groups were performed using non-parametric Mann–Whitney U tests. Comparisons involving multiple groups were analyzed using two-way ANOVA followed by Šídák’s multiple comparisons test for parametric data, or by the Kruskal–Wallis test for nonparametric data. Simple linear regression analysis was used to assess correlations and assay calibration. ROC was performed to evaluate assay sensitivity and specificity. Detailed information on statistical tests, sample sizes, and significance levels is provided in the corresponding figure legends.

## Data Availability

All data supporting the findings of this study are included in the article and its supplementary information files. Additional raw data and analysis materials are available from the corresponding author upon reasonable request.

## Results

### Plasma cells synthesize JCHAIN independent of IgH isotype

The cellular source of mIgA and dIgA in mice and humans remain incompletely defined. A prevalent model postulates that plasma cells in systemic lymphoid tissues, such as the bone marrow, account for the substantial amounts of the mIgA in circulation (20,21) while dIgA selectively arises from dedicated plasma cells in mucosal tissue (19). This binary model predicts the existence of functionally specialized plasma cell subsets secreting either mIgA or dIgA. However, individual IgA plasma cells may produce both molecular forms, thereby contributing simultaneously to monomeric and dimeric IgA pools (Fig. 1A). To assess the molecular forms of mucosal and circulating IgA, we performed a non-reducing Western blot of IgA in saliva, feces, and serum of wild-type mice (Fig. 1B). As expected, saliva and fecal samples showed a high molecular weight form corresponding to sIgA. This represents IgA dimers and multimers complexed with JCHAIN and SC, reflecting PIGR-mediated transport across epithelial barriers. In contrast, serum IgA was dominated by bands corresponding to mIgA, in agreement with the established enrichment of mIgA in the circulation.

**Figure 1.**
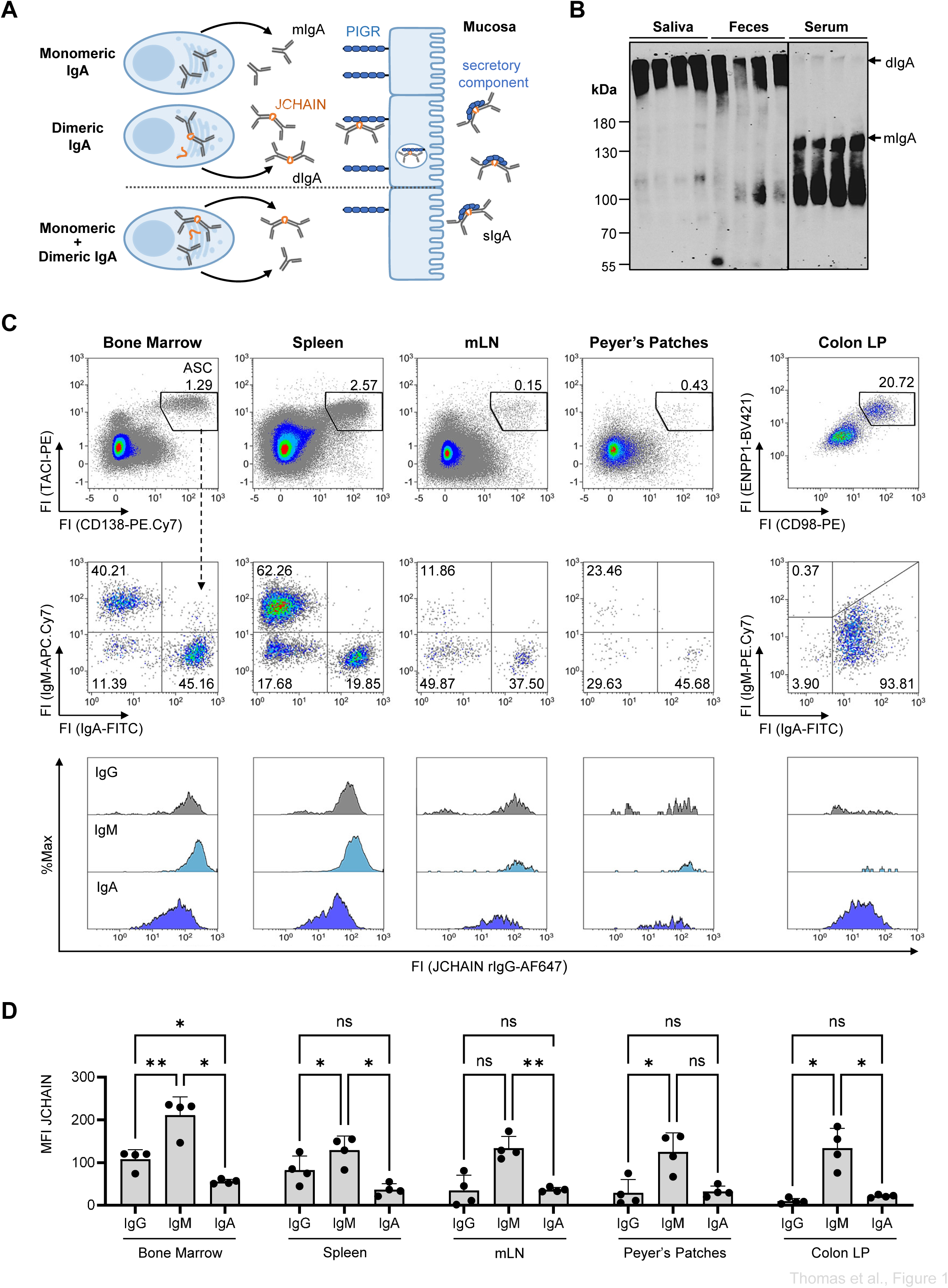
JCHAIN protein abundance in ASCs across tissues. **A.** Schematic representation of potential cellular sources and epithelial transport pathways of monomeric IgA (mIgA), dimeric IgA (dIgA), and secretory IgA. **B.** Non-reducing Western blot analysis of saliva, feces and serum from C57BL/6 mice, detected with HRP-conjugated anti-mouse IgA. Bands corresponding to sIgA (∼384 kDa), dIgA (∼ 318 kDa) and mIgA (∼150kDa) are indicated on the right. Molecular mass standards (kDa) are indicated on the left. Each lane represents one mouse. **C.** Representative flow cytometric analysis of intracellular JCHAIN abundances in ASCs from the indicated tissues. Numbers indicate the percentage of cells within the respective gate. **D.** Quantification of JCHAIN abundances, shown as median fluorescence intensity (MFI), in ASC populations gated as in C. Statistical analysis was performed using a two-way ANOVA with Šídák’s multiple comparisons test. Data are shown as mean +/- SD, n=4, data represent 2 independent experiments. Ns, non-significant, *p < 0.05 and **p < 0.01. PIGR = polymeric immunoglobulin receptor, mLN = mesenteric lymph nodes, Colon LP = colon lamina propria

Given the essential role for JCHAIN in the formation of canonical IgA dimers, JCHAIN-negative IgA plasma cells have been proposed as a source of mIgA (reviewed in Castro and Flajnik, 2014). To challenge this notion, we determined whether antibody-secreting cells (ASC) lacking intracellular JCHAIN exist, representing a potential source of mIgA. Intracellular JCHAIN abundance was determined in the ASC compartments, comprising plasmablasts and mature plasma cells. ASC were identified by CD138/TACI expression in the bone marrow, spleen, mesenteric lymph nodes (mLN) and Peyer’s patches (PP) and were further stratified by surface Ig expression (22) (complete gating strategy depicted in Fig. S1). CD98 and ENPP1 were used as alternative surface markers for colon lamina propria (cLP) plasma cells due to loss of CD138 and TACI from the cell surface by enzymatic digest during the isolation procedure (23–26), in combination with intracellular staining against IgA and IgM heavy chains. In unimmunized wild-type mice, the IgA- and IgM-double negative ASCs predominantly express IgG antibodies (22,27) and therefore, we refer to this population as IgG ASCs. Unlike predicted by the binary model, flow cytometric analysis demonstrated that nearly all ASCs in the analyzed tissues express JCHAIN protein, regardless of Ig heavy chain isotype, although a small fraction of JCHAIN-negative events was detected within the IgG ASC population (Fig. 1C). However, JCHAIN abundance varied between ASC subsets: IgM-positive ASCs displayed the highest JCHAIN staining intensities, IgG ASCs exhibited intermediate JCHAIN abundance and IgA-positive ASCs showed surprisingly the lowest intensities among the ASC subsets analyzed (Fig. 1C, D). Nevertheless, JCHAIN expression remained detectable above background in IgA-producing ASCs from all analyzed tissues (Fig. S2A), including the bone marrow, suggesting that the presence of mIgA cannot be explained by a population of IgA ASCs completely lacking intracellular JCHAIN. Moreover, given the prediction that bone marrow cells synthesize and secrete mIgA, it was striking to see that JCHAIN staining intensities were even elevated in bone marrow ASCs compared with their counterparts in the other analyzed tissues (Fig. 1D). Hence, our data do not provide evidence for a discrete JCHAIN-negative ASC population accounting for mIgA production.

### Recombinant soluble PIGR selectively detects dimeric IgA

We did not identify a discrete JCHAIN-negative ASC population. Notwithstanding, JCHAIN expression alone is not sufficient for dIgA synthesis, as additional factors support its formation (18). To develop a direct method for dIgA detection, we employed the PIGR ectodomain, also known as the secretory component. The PIGR ectodomain consists of five Ig-like domains which mediate the covalent binding of the PIGR to JCHAIN and the IgA heavy chains (28). We produced the murine PIGR ectodomain as a soluble secreted protein in 293F cells, incorporating C-terminal 3xFLAG- and Strep-tags for detection and purification (Fig. 2A, Fig. S2B). We then tested binding of the purified sPIGR to 293T cells transfected with expression plasmids for mouse JCHAIN, IgA heavy chains or combinations thereof (Fig. 2B). While the anti-JCHAIN antibody stained transfected JCHAIN in the absence and presence of co-transfected IgA heavy chains, sPIGR binding was only observed when both IgA and JCHAIN were co-expressed. However, the sPIGR signal intensities were modest and restricted to cells with highest IgA abundances, possibly reflecting inefficient intracellular assembly in the heterologous expression system. Nevertheless, these results provided initial evidence that the sPIGR reagent specifically detects JCHAIN-containing multimeric IgA.

**Figure 2.**
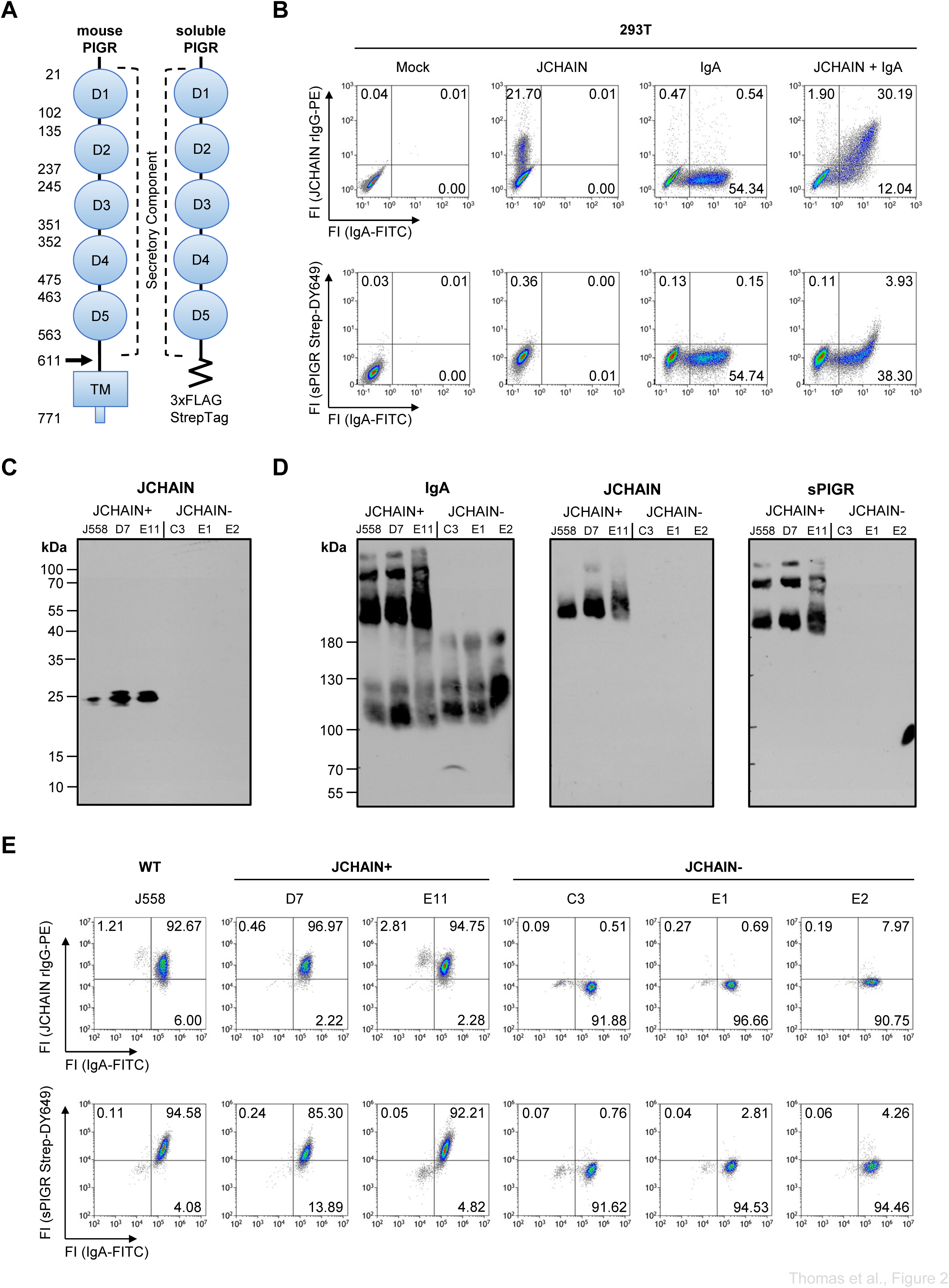
Specificity of recombinant sPIGR in flow cytometry and Western blot analyses. **A.** Schematic representation of the murine polymeric Ig receptor (PIGR) and the recombinant sPIGR construct with C-terminal tags. Numbers indicate amino acid positions in reference sequence (NP_035212.2), and the arrows denote the proteolytic cleavage site. TM, transmembrane domain **B.** Flow cytometric analysis of fixed and permeabilized 293T cells transiently transfected with expression plasmids encoding mouse JCHAIN, a murine IgA heavy chain (derived from J558 cell line), or both plasmids in combination. Intracellular JCHAIN and IgA heavy chain were detected as indicated. Binding of recombinant sPIGR was detected by subsequent staining with Strep-Tactin XT-Dy649. Data represent 2 independent experiments. **C.** J558 mouse plasmacytoma cells were transfected with Cas9/RNP-complexes targeting the mouse *Jchain* locus. Single cell clones were sorted, expanded and cell supernatants were analyzed for JCHAIN expression (∼18kDa) by Western blot under reducing conditions. **D.** Non-reducing Western blot analysis of culture supernatants from the J558-derived JCHAIN-positive and -deficient clones. The blot was sequentially probed for IgA, JCHAIN and soluble PIGR-binding. Bound sPIGR was detected with HRP-conjugated anti-FLAG antibody. **E.** Flow cytometric analysis of J558-derived JCHAIN-positive and -deficient clones. Cells were fixed, permeabilized and stained with anti-JCHAIN antibody followed by PE-conjugated anti-rabbit IgG, together with anti-IgA-FITC detection of IgA heavy chain expression (upper panel). In the lower panels, cells were stained intracellularly with anti-IgA-FITC antibody and incubated with recombinant sPIGR, followed by detection with Strep-Tactin XT-Dy649. Data represent 2 independent experiments.

We further assessed the specificity of the sPIGR in the murine IgA-secreting plasmacytoma cell line J558 (29) and in JCHAIN-deficient J558 clones generated by CRISPR/Cas9 gene editing. Western blot analyses under reducing conditions of the parental J558 line and selected clones confirmed successful *Jchain* targeting, with no detectable JCHAIN bands in the supernatants of the knock-out clones (Fig. 2C). Western blot analysis under non-reducing conditions confirmed that the J558 cells secrete both mIgA and dIgA (Fig. 2D). In the cell culture supernatants from the parental J558 cells and JCHAIN-expressing clones, high-molecular weight IgA complexes were detectable that also contained JCHAIN. In contrast, JCHAIN-deficient J558 clones lacked dIgA and higher-order complexes and secreted only mIgA. Notably, sPIGR binding was restricted to the JCHAIN-containing dIgA or higher-order complexes, establishing this reagent as a specific probe for multimeric IgA in Western blot assays (Fig. 2D).

Next, we compared the intracellular sPIGR binding by flow cytometry in the parental J558 line and selected clones with or without *jchain* deletions (Fig. 2E). The parental J558 line and JCHAIN-producing clones showed specific sPIGR binding, whereas sPIGR binding was abolished in JCHAIN-deficient J558 clones, despite comparable detection of intracellular IgA. Thus, sPIGR binding required the combined presence of IgA heavy chains and JCHAIN, indicating that sPIGR recognizes JCHAIN-associated IgA complexes rather than IgA or JCHAIN alone. In addition, JCHAIN abundance has no influence on IgA abundance in this system. Together, these data establish sPIGR as a specific reagent for distinguishing mIgA from multimeric IgA in Western blot and flow cytometric analyses.

### Soluble PIGR reveals universal dIgA formation by IgA plasma cells

With sPIGR validated as a reagent for detecting dIgA, we next asked whether IgA plasma cells differ in their capacity to form PIGR-reactive IgA despite expressing JCHAIN. The identification of an IgA plasma cell subset failing to bind the sPIGR could represent a dedicated cellular source of circulating mIgA. We analyzed ASCs in the systemic and mucosal lymphoid tissues of wild-type mice, with intracellular staining for the Ig isotype and detection of sPIGR binding (Fig. 3A). Soluble PIGR binding was not detected in IgG ASCs, whereas IgM ASCs showed only a modest signal shift. In contrast, IgA ASCs displayed strong sPIGR binding across all analyzed tissues. Similar to the pattern observed for JCHAIN, bone marrow ASCs exhibited the highest sPIGR binding intensities, whereas ASCs in the cLP showed the lowest signals (Fig. 3B). When sPIGR binding was normalized to intracellular IgA fluorescence, IgA⁺ ASCs from the bone marrow and cLP showed comparable relative sPIGR binding (Fig. 3C). Given the demonstrated specificity of sPIGR for dIgA, the absence of a discrete sPIGR-negative IgA⁺ ASC population argues against the existence of a dedicated subset secreting only mIgA. Instead, these data indicate that all IgA ASCs in both systemic and mucosal tissues are capable of forming dIgA.

**Figure 3.**
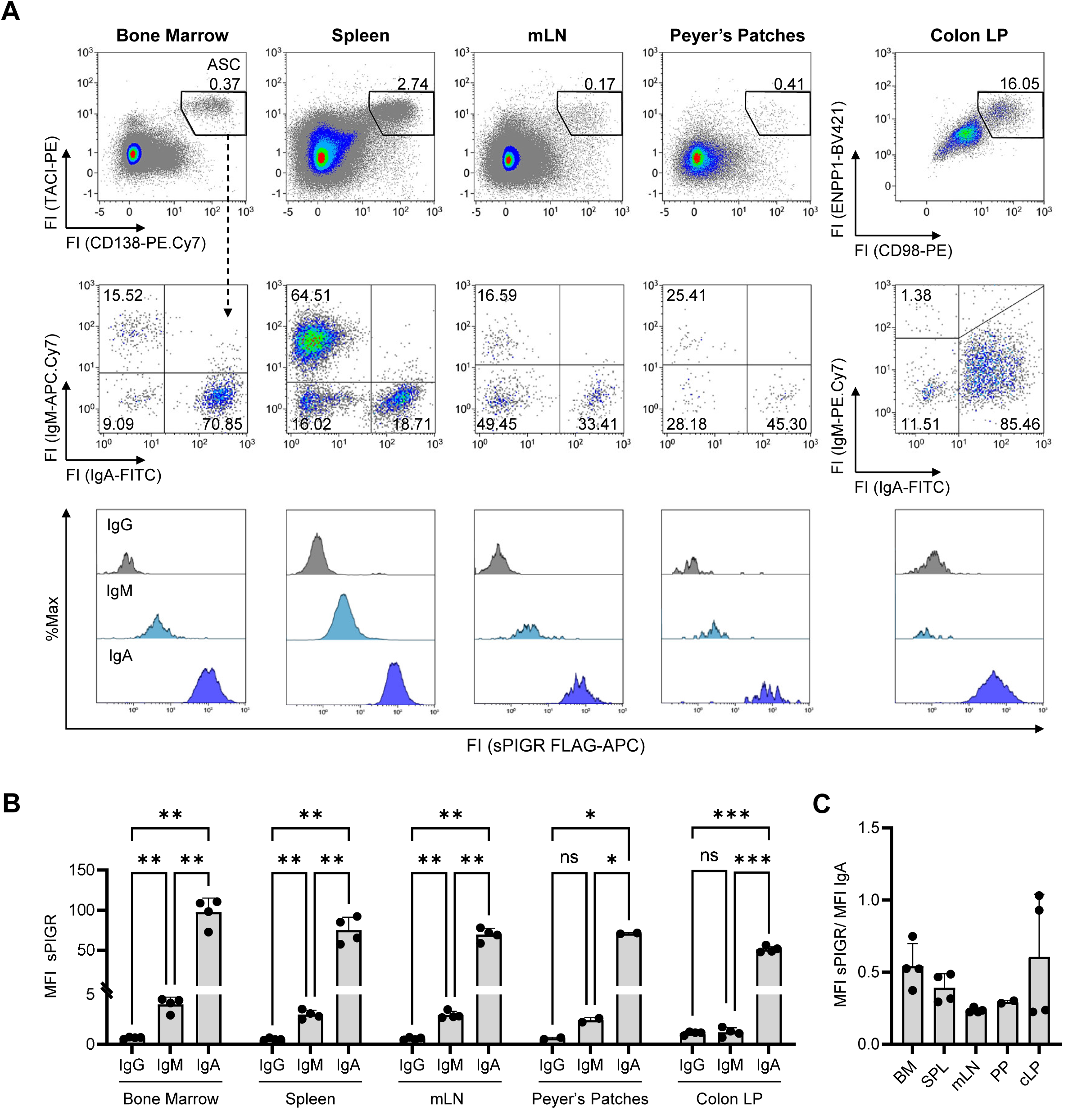
Flow cytometric analysis of sPIGR binding in ASCs across tissues. **A.** Single cell suspensions from murine bone marrow, spleen, mesenteric lymph nodes (mLN), Peyer’s patches and colonic lamina propria (Colon LP) were surface-stained for TACI and CD138, followed by intracellular staining for IgA and IgM. Intracellular binding of sPIGR was detected using anti-FLAG-APC. Representative histograms show sPIGR/anti-FLAG-APC fluorescence across IgH-gated ASCs subsets. **B.** Quantification of sPIGR binding via median fluorescence intensities (MFI) across tissues and ASC isotypes. Statistical analysis was performed using a two-way ANOVA with Šídák’s multiple comparisons test. Data are shown as mean ± SD; n=4, data represent 2 independent experiments. Ns, non-significant, *p < 0.05, **p < 0.01 and ***p < 0.001. **C.** Ratio of sPIGR/ IgA MFIs in IgA ASCs across tissues. Data are shown as mean ± SD; n=4. BM= bone marrow, SPL= spleen, mLN= mesenteric lymph nodes, PP= Peyer’s patches and cLP= colonic lamina propria

### Bone marrow plasma cells secrete both monomeric and dimeric IgA

Having shown JCHAIN presence and sPIGR binding in all IgA plasma cells, the source of mIgA remained enigmatic, particularly because a flow cytometric reagent that selectively identify mIgA is not available. PIGR binding does not exclude parallel synthesis and secretion of mIgA and dIgA. To determine whether IgA plasma cells secrete mIgA and dIgA simultaneously, we established a PIGR sandwich ELISA. Total IgA was first captured with anti-IgA antibodies and detected either with polyclonal anti-IgA to measure total IgA or with sPIGR to selectively detect dIgA and higher-order IgA complexes. Assay performance was validated using recombinant mouse mIgA and dIgA generated by co-transfection of 293F cells with Igα and Igκ expression vectors with or without JCHAIN, as previously described (30). Defined dIgA:mIgA ratios were prepared while maintaining a constant total IgA concentration. The sPIGR signal showed a strong linear correlation with the dIgA content, confirming that the assay quantitatively reports the dIgA fraction in mixed IgA samples (Fig. 4A).

**Figure 4.**
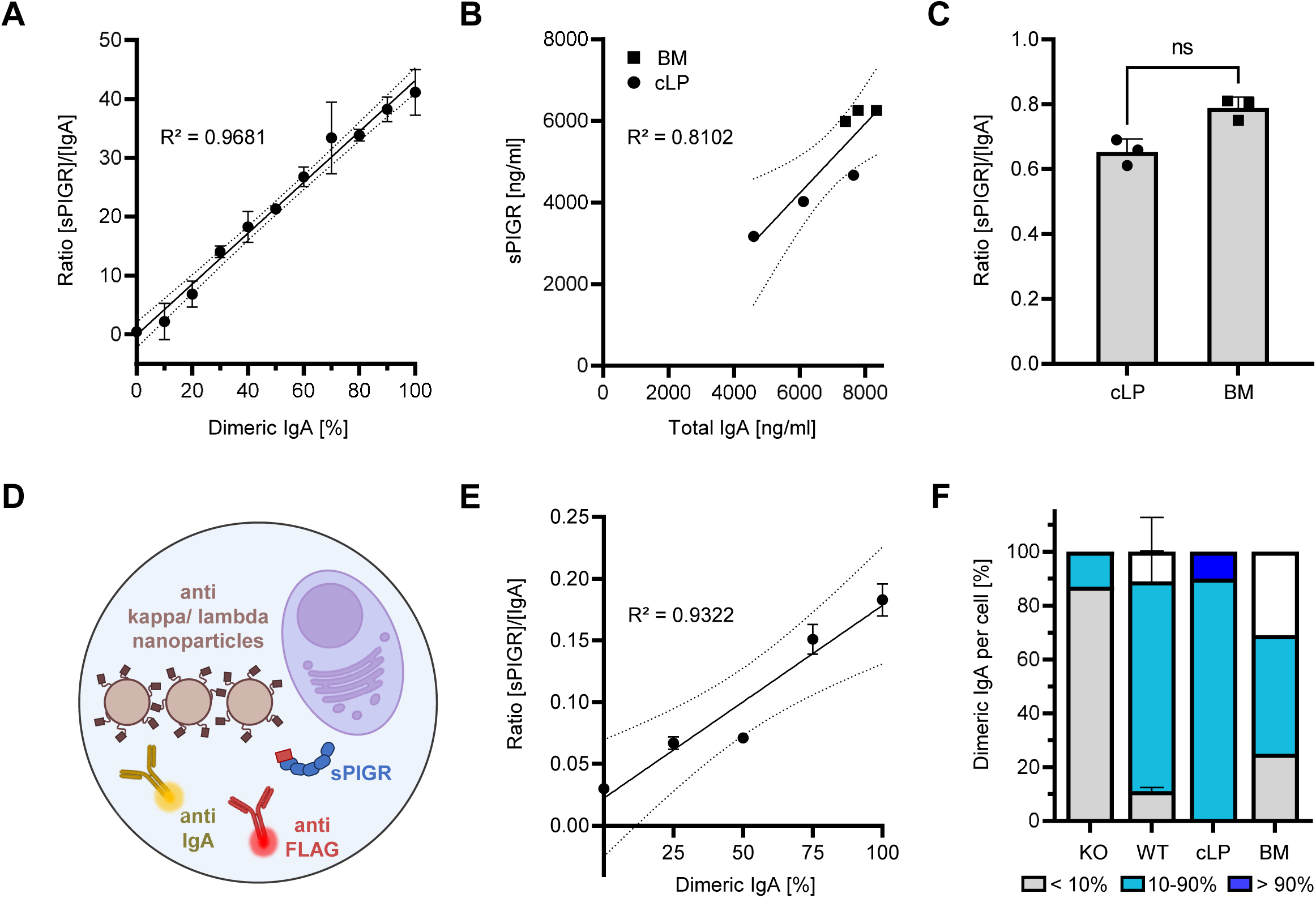
Bulk and single-cell quantification of dimeric IgA secretion from ASCs. **A.** Calibration of the sPIGR-based ELISA using defined mixtures of recombinant monomeric and dimeric IgA. IgA mixtures were analyzed by ELISA using anti-IgA antibodies and recombinant sPIGR. The ratio of sPIGR binding to total IgA abundance is plotted against the percentage of dimeric IgA in the input mixture. Data were fitted by linear regression with the corresponding 95% confidence intervals. **B.** IgA-producing ASCs (IgA^+^ TACI^+^ CD138^+^) were sorted from the bone marrow (BM) and colon lamina propria (cLP) and cultured for 24h in R10 medium supplemented with APRIL. Supernatants were analyzed by IgA ELISA and sPIGR-based ELISA. Each dot represents pooled cells of each four individual C57Bl/6 mice; line indicates simple linear regression with the corresponding 95% confidence intervals. **C.** Ratio of sPIGR binding to total IgA abundance in culture supernatants from BM- and cLP-derived IgA ASCs cultured for 24 hours in R10 medium supplemented with APRIL. Statistical significance was assessed using the Mann-Whitney U-test; ns, non-significant. **D.** Schematic representation of the DropMap experimental setup. Magnetic nanobeads coated with anti-Igκ or anti-Igλ antibodies were embedded in microfluidic droplets together with single cells and detection reagents (anti-IgA-FITC, sPIGR, anti-FLAG-AF647). Antibody secretion is visualized by fluorescence relocation onto magnetic particles and quantified. **E.** Calibration of DropMap analysis using defined mixtures of recombinant dIgA and mIgA. The ratio of the sPIGR-dependent anti-FLAG-AF647 signal to the IgA-FITC signal is plotted against the percentage of dimeric IgA in the input mixture. line indicates simple linear regression with the corresponding 95% confidence intervals. **F.** Parental J558 (WT) or JCHAIN-deficient J558 cells (KO) (2 experiments), as well as sorted IgA-producing ASCs from colon lamina propria (cLP) or bone marrow (BM) pooled from three C57Bl/6 mice were analyzed by DropMap. Secreted dIgA detected by sPIGR binding was quantified relative to total IgA. Bars show the frequencies of cells classified as secreting <10% IgA dimer, 10-90% IgA dimer or >90% IgA dimer.

We next applied this ELISA to supernatants from bulk-sorted primary IgA-positive ASCs isolated from bone marrow and cLP. Bone marrow IgA ASCs secreted higher amounts of total IgA compared to cLP ASCs, consistent with recent evidence that gut-resident plasma cells display comparatively low IgA secretion (31). Importantly, IgA secreted by cLP ASC, but also IgA secreted by bone marrow ASC, showed clear sPIGR binding, indicating that both tissue ASC populations secrete dIgA (Fig. 4B). Unexpectedly, the relative dIgA fraction among total secreted IgA was higher in bone marrow ASC cultures than in cLP ASC cultures, although this difference did not reach statistical significance (Fig. 4C). Thus, rather than supporting a model in which bone marrow IgA ASCs predominantly or exclusively secrete mIgA, this finding demonstrates clearly that bone marrow IgA ASCs are a source of dIgA. This result predicted simultaneous secretion of mIgA and dIgA by the same cell.

To test this prediction, we established a quantitative single-cell assay based on single droplet analysis (32) (Fig. 4D). This approach enabled simultaneous fluorescent detection of total IgA and sPIGR-reactive dIgA secreted by individual ASCs in droplets (DropMap assay). Assay performance was calibrated with defined recombinant mIgA/dIgA mixtures, which showed a proportional relationship between dIgA content and sPIGR signal in droplets (Fig. 4E). Within the limits of the assay’s specificity and sensitivity (Fig. S2C), JCHAIN-deficient J558 clones secreted only mIgA, confirming that the DropMap sPIGR signal depends on JCHAIN-mediated IgA multimerization (Fig. S2C Fig. S2D). Most J558 cells in droplets secreted between 10-90% dIgA and only a minority secreted dominantly mIgA or dIgA (Fig. 4F, Fig. S2D), which is consistent with our Western blot analyses (Fig. 2D).

We then encapsulated primary IgA ASCs from bone marrow and cLP for DropMap analysis. Consistent with the bulk secretion data, the majority of ASCs from both tissues secreted detectable dIgA as a fraction of total IgA. Importantly, the majority of IgA ASCs from both tissues were classified as dual secretors (10–90% sPIGR-bound IgA), rather than falling into the mIgA-only or dIgA-only categories, which are defined as having less than 10% of the reciprocal IgA form (see Fig. 4F and Fig. S2D). Because total IgA and sPIGR-binding IgA were measured in the same droplets, these data indicate that the majority of individual IgA ASCs concomitantly secrete both mIgA and dIgA. Thus, secretion of dIgA and mIgA is not restricted to IgA ASCs in mucosal or systemic lymphoid tissues, respectively. Instead, the capacity to secrete both IgA molecular forms in parallel appears to be a general feature of IgA ASCs.

## Discussion

Secretory IgA is thought to be produced by mucosal IgA plasma cells that assemble JCHAIN-containing dIgA locally (19). In contrast, the origin of circulating mIgA is often traced back to plasma cell subsets that are anatomically or functionally distinct from mucosal plasma cells. Such distinctions could result from alterations in JCHAIN expression or other factors influencing dIgA assembly. In this study, we tested this model by combining intracellular JCHAIN detection (25,33) and direct detection of dIgA with a recombinant sPIGR probe. The use of the sPIGR was validated for Western blot, flow cytometry, ELISA and DropMap analysis (32), establishing this reagent as a versatile tool for detecting native and recombinant polymeric IgA. Using JCHAIN detection and the sPIGR, we found that all IgA plasma cells from murine mucosal and systemic lymphoid tissues uniformly express JCHAIN, consequently bind sPIGR, and secrete PIGR-reactive IgA. Moreover, single-cell DropMap analysis (32) showed that most individual murine IgA ASCs isolated from bone marrow and cLP co-secrete mIgA and dIgA rather than producing either molecular form exclusively. These findings argue against a binary distinction of mIgA-only and dIgA-only IgA plasma cell populations. Instead, they support a model in which dIgA assembly is a common feature of all IgA plasma cells, whereas the relative output of mIgA and dIgA is regulated quantitatively within individual cells and tissue environments. This regulation may differ across species, depend on the general immune status, age or other factors and does not exclude a putative expansion of mIgA secreting cells under certain conditions. The *Jchain* transcription and protein abundance can be uncoupled, as post-transcriptional regulation or degradation of unassembled JCHAIN in ASCs lacking polymerizing Ig isotypes occurs (34). Our findings clearly reveal that there are no JCHAIN negative plasma cells. Transcriptomic analyses and models of plasma cell differentiation (38–40) also support the view that *Jchain* transcription is induced as part of the ASC transcription program, whereas earlier work has raised the possibility of JCHAIN-negative plasma cells, particularly among mIgA-secreting cells and some IgG-producing ASCs (5,6). To our knowledge, the present study provides the first flow cytometric analysis that detects JCHAIN protein in mouse ASC subsets from systemic and mucosal tissues. We show that JCHAIN protein expression is a general feature in all ASCs, including IgA, IgM, and IgG ASCs. Thus, our data argue against selective post-transcriptional elimination or degradation of JCHAIN in ASCs that express non-polymerizing Ig isotypes. Instead, they are consistent with transcriptional models in which JCHAIN expression is generally linked to plasma-cell differentiation by relief of PAX5-dependent repression during terminal differentiation (41). The surprising finding that JCHAIN abundance was highest in IgG ASCs further raises the possibility that JCHAIN may have functions in ASCs beyond Ig polymerization. This remains speculative, but is conceptually supported by recent evolutionary work identifying JCHAIN as an evolutionarily co-opted CXCL chemokine family member (42). One possible explanation for the apparent discrepancy between our findings and earlier reports of JCHAIN-negative ASCs is that the previous detection of JCHAIN was technically challenging and required denaturing pretreatments to unmask epitopes. This could lead to an underestimation of the number of JCHAIN-expressing ASCs (5). A revised analysis of IgG ASC in *jchain*^-/-^ mice (43,44) might unravel specific roles of JCHAIN beyond Ig polymerization. To specifically identify dIgA-producing ASCs, we validated the use of sPIGR as a detector of JCHAIN-containing dIgA in ELISA, DropMap, Western blot and flow cytometry. Of note, we produced a defined, recombinant sPIGR, while purified SC from colostrum has previously been used to distinguish two types of human IgA-producing cells, containing monomeric versus dimeric IgA (19). IgA-positive cells binding SC in the mucosa were defined as dIgA producing cells, while SC-negative cells found in other glands were defined as mIgA producing cells (19). This study, however, did not analyze bone marrow, spleen or other lymphoid tissues and was performed in humans. Therefore, our experiments extend these findings.

The binding of sPIGR to virtually all mouse IgA ASCs by flow cytometry provides evidence for their competence to assemble dIgA. However, this alone is insufficient to determine the relative proportions of mIgA and dIgA secreted by individual ASCs. We addressed this important objection first by quantifying bulk secretion of total IgA and dIgA from cLP and bone marrow IgA plasma cells in the presence of APRIL. Both populations secreted substantial amounts of sPIGR-binding IgA, demonstrating that IgA ASCs from systemic and mucosal tissues release dIgA. Notably, bone marrow ASC culture supernatants contained higher total IgA amounts from the same number of seeded cells and also a higher relative fraction of dIgA compared with their cLP counterparts, a finding that corresponds with a recent report (35).

Earlier work has already demonstrated the capacity of human IgA ASCs from various tissues to form and secrete dIgA, suggesting predominant co-secretion of both mIgA and dIgA from individual ASCs (45). Our data are in support of these findings. Moreover, clonal profiling of human IgA1 revealed shared clones among the mIgA1 and dIgA1 fractions (46). Apparent discrepancies with reports of distinct clonal repertoires between serum mIgA and breast-milk dIgA (47) may reflect anatomical compartmentalization rather than mutually exclusive mIgA-or dIgA-producing ASC lineages. Individual IgA ASCs may be capable of producing both molecular forms, while tissue localization, PIGR-mediated epithelial transport, and rapid clearance of polymeric IgA from the circulation via the biliary route (7) determine which IgA species are ultimately detected in serum or other body fluids.

Our proposal integrates the functions of systemic and mucosal IgA. Flexible production of mIgA and dIgA may be relevant for microbiota containment and systemic defense against mucosal pathogens. However, the triggers that determine the mIgA:dIgA output ratio of individual ASCs remain unknown. Potential regulators include the relative abundance of IgA heavy chains, light chains, and JCHAIN, ER-folding and quality-control pathways including chaperones such as MZB1 (18), tissue-derived signals or inflammatory cues. Defining how these factors quantitatively tune IgA polymerization will be important for understanding how systemic and mucosal IgA functions are coordinated during steady-state immunity, vaccination, and infection. To conclude, our findings demonstrate that dIgA competence is shared among all mouse IgA plasma cells rather than confined to a discrete mucosal plasma cell subset. We propose that the secreted dIgA:mIgA ratio is regulated within the IgA plasma cell lineage by cell-intrinsic factors, with bone marrow IgA plasma cells representing a clear systemic source of dIgA.

## Acknowledgments

This work was supported by grants from the Deutsche Forschungsgemeinschaft (DFG) GK2599 (to D.M., H.M.J), Mi939/6-1, project 503852185, to D.M., Interdisciplinary Clinical Research Center (IZKF) Erlangen (ER-Stress und Sialylierung in Plasmazellen), to D.M., Project-ID 506620580-Emmy Noether Grant and Project-ID 390873048 - EXC2151 to T.R. We thank the Core Unit Cell Sorting of the Friedrich-Alexander-Universität Erlangen-Nürnberg and FACS Core Facility of Aarhus University for excellent technical assistance.

## Declaration of Generative AI and AI-assisted technologies in the writing process

During the preparation of this work, the authors used ChatGPT (OpenAI, GPT-5.5, DeepL Write) to improve the clarity and readability of individual sentences. After using this tool, the authors reviewed and edited the content as needed and take full responsibility for the content of the published article.

## Author Contributions

S.R.S., D.M. and H.-M.J.: Conceptualization. S.R.S., J.T. and D.M.: Methodology and experimental design. K.E.: Investigation, formal analysis, and data visualization for DropMap experiments. J.W., E.R. and W.X.: Investigation. T.R.: Resources. W.S., D.M. and H.-M.J.: Supervision. J.T., D.M. and S.R.S.: Writing – original draft. All authors: Writing – review & editing.

## Abbreviations

ASC: antibody-secreting cells
BM: bone marrow
cLP: colon lamina propria
dIgA: dimeric IgA
ER: endoplasmic reticulum
FL: fluorescence
FSc: Forward Scatter
GFP: green fluorescent protein
Ig: immunoglobulin
JCHAIN: joining chain
KO: knock-out
MFI: median fluorescence intensity
mIgA: monomeric IgA
MZB1: marginal zone B and B 1 cell specific protein 1
mLN: mesenteric lymph nodes
PIGR: poly Ig receptor
PP: Peyer’s patches
SC: secretory component
sIgA: secretory IgA
sPIGR: soluble PIGR
SPL: spleen
SSc: Side scatter
WT: wildtype.

---

**Figure S1 Gating strategy for identification of ASCs**

Flow cytometric gating strategy used to identify TACI⁺ CD138⁺ ASCs and IgA⁺, IgM⁺, and IgA⁻IgM⁻ (IgG) ASC subsets, shown representatively for the bone marrow.

**Figure S2 Detection of dimeric IgA in mouse plasma cells and single-cell secretion assay using recombinant sPIGR**

**A.** Flow cytometric analysis of ASCs (TACI^+^CD138^+^) from murine bone marrow, spleen, mesenteric lymph nodes (mLN), Peyer’s patches and colonic lamina propria (Colon LP). Upper panels show staining with Alexa Fluor 647-conjugated anti-rabbit IgG secondary antibody alone or following incubation with rabbit anti-JCHAIN monoclonal antibody. Lower panels show staining with recombinant soluble PIGR (sPIGR) or an irrelevant FLAG-tagged control protein, followed by detection with APC-conjugated mouse anti-FLAG antibody.

**B.** Coomassie-stained SDS gel of purified sPIGR (∼ 72.5 kDa) separated under reducing (R) and non-reducing (NR) conditions.

**C.** Receiver operator characteristic (ROC) of the sPIGR-based DropMap assay for the discrimination of J558 WT and JCHAIN-deficient J558.

**D.** DropMap analysis of parental J558 (WT) and JCHAIN-deficient J558 (KO) (2 experiments) and sorted IgA⁺ ASCs from colonic lamina propria (cLP) and bone marrow (BM) pooled from three C57Bl/6 mice. Secreted dimeric IgA was detected by sPIGR binding followed by anti-FLAG-AF647 and quantified relative to total IgA detected with anti-IgA-FITC. Each dot represents one cell; red lines indicate the mean. Statistical analysis was performed by Kruskal–Wallis ANOVA. Ns, non-significant, ***p < 0.001 and ****p < 0.0001.

