## Supplementary figures and images for "IgA plasma cells co-secrete monomeric and dimeric IgA"

### Supplemental Figures

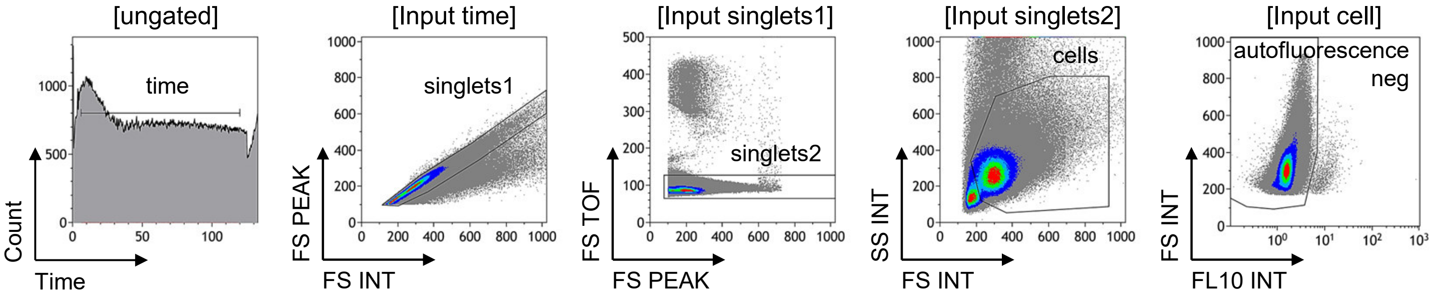


Thomas et al., Figure S1

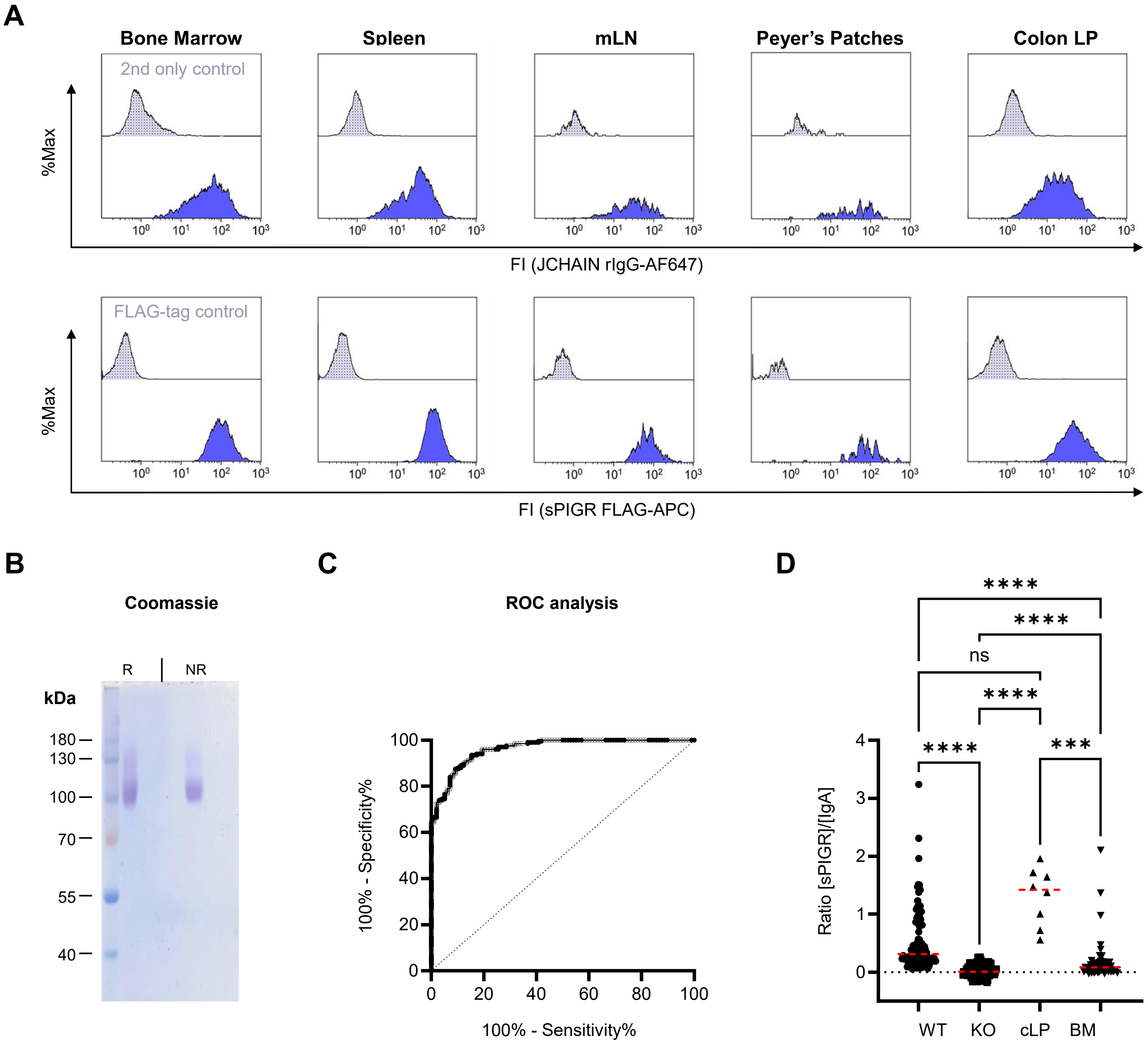


Thomas et al., Figure S2
